## Supplementary material for "Reassessment of the basis of cell size control based on analysis of cell-to-cell variability": Supplermetary Figures and Tables

### MATERIALS AND METHODS

#### Image acquisition and analysis

All images analysed here are taken from our previous publication [1]. High resolution images were used to obtain cell segmentation at different resolutions by binning pixels (Figs. 2, 3, S1). This allowed us to study the effect of the segmentation error. In these cases, we applied the following new semi-automated algorithm to the images acquired at the best z-plane. We first drew manually an approximate cell axis that connected the two cell tips (A and B, see Fig. S7A). The AB distance was a first estimation *L* of the cell length. A first estimation of the cell radius was calculated as follows. From the middle point M of the AB segment, we derived an intensity profile along the direction orthogonal to the axis (towards both lateral borders of the cell, red dashed line in Fig. S7A). The steepest gradient of this signal identified the border of the cell, i.e. the cell radius. We repeated this gradient procedure using neighbouring points of M (at a distance $k \cdot dx$, with k=-3, -2, … , +2, +3, and where dx is the pixel size). The average of all these values gave our first estimation of the radius. We then shortened the segment AB by this amount from both ends. This defined the new segment A’B’. We divided this segment into n equal parts ($n=0.5 \cdot L/dx$) by placing n-1 internal points. The gradient procedure we used for the middle point M was then applied to all these points and to the two extremal points A’ and B’. This identified the lateral borders of the cells. The symmetry axis of the resulting lateral borders was taken to be the new symmetry axis of the cell (segment CD). From both extreme points C and D, we derived the intensity profile along the different radial directions (from 0 to 180 degrees, steps of 2 degrees, purple dashed line in Fig. S7B). Ideally, the distance of the cell border to the CD segment should always have been equal to R for every point from C to D. However, cell irregularity and imaging defects altered this width. Therefore, after calculating the width from a point $x_i$, if the difference between this width and the width from the previous point $x_{i-1}$ was bigger than a given threshold (0.1 μm), we set the width equal to the width from $x_{i-1}$. The final result was a set of points that described the cell border. From these points we derived the cell length as the maximal distance between two points of the set (Fig. S7C). The cell radius was defined as the average of its value at different positions along the CD segment, i.e. excluding the two cell tips (Fig. S7C). Surface area and volume were then calculated using simple geometrical equations. As showed by Eq. 2 of the main text, segmentation error (i.e. high image resolution) is not critical in case of calculation of the division-birth slope based on length measurements (Fig. 1, 4, S2, S4B and S5). In these cases, phase contrast images were first analysed by a deep neural network machine learning algorithm [2]: this methodology generated binary images for feature (outline/cytoplasm) identification. These contours were then used for traditional gradient segmentation in Morphometrics [3]. This procedure is fully described in [1]. Strains used in the experiments are listed in Table S2.

**Equations for the pure sizer model**

Cells were assumed to have a perfect cylindrical shape with hemispherical ends. Surface area and volume were calculated using values of the cell length and of the cell radius: $A = 2\pi RL$ and $V = \pi R^2 L - 2\pi R^3/3$. We initially assumed no variation of the cell width during a single cell cycle, although later this assumption was relaxed.

**Notations:** In the following: $\mu$ is the relative error of the cell in sensing size, $\rho$ is the relative cell to cell variability of the radius (so that the true radius of a cell is $R(1+\rho)$), $\alpha$ is the relative error at division due to asymmetric misplacement of the septum (so that cell does not divide symmetrically in two halves but into $(1+\alpha)/2$ and $(1-\alpha)/2$ fractions), and $\varepsilon$ is the experimental error (in μm) in measuring distances. All these quantities are assumed to have a Gaussian distribution with mean zero. The following calculations will then be extended to negative values of the geometrical quantities but where the probability weight of these tails is too small to have a significant impact on the results. We also need different copies of some random variables in order to describe the variation at different moments of the cell cycle: for instance, depending on whether we are considering birth or division, the relative error of the cell in sensing size at division must be described by $\mu_b$ or $\mu_d$, respectively. These are independent and identically distributed random variables. The same holds for the error in our measurements ($\varepsilon_b$ for the length at birth, $\varepsilon_d$ for the length at division and $\varepsilon_r$ for the cell radius).

**Cell length, area and volume for the case of surface area sensing:** By stating that a cell is sensing surface area, this means that division occurs at a given area $A$ (plus the error $A\mu_d$ due to the cellular error in size sensing). Therefore, the real division length ($L_d$, not affected by our error in its measurement) of a cell with real radius $R(1+\rho)$ is the length such that the resulting cell area is equal to the target value corrected by the error due to the imperfect cell sensing, i.e. $2\pi R(1+\rho)L_d = A(1+\mu_d)$. By adding the error $\varepsilon_d$ of our measurement, we have the corresponding measured quantity $L_d^*$:

$$L_d^* = \frac{A(1+\mu_d)}{2\pi R(1+\rho)} + \varepsilon_d. \qquad \text{(S1)}$$

Following similar reasoning, we derive the measured radius $R^* = R(1+\rho) + \varepsilon_r$. Because of the experimental errors, the expression of the measured area at division does not coincide with the theoretical value corrected by the sensing error, i.e. $A(1+\mu_d)$. By definition, the measured area at division is

$$A_d^* = 2\pi R^* L_d^* = 2\pi[R(1+\rho) + \varepsilon_r]\left[\frac{A(1+\mu_d)}{2\pi R(1+\rho)} + \varepsilon_d\right], \tag{S2}$$

while the measured volume at division is

$$V_d^* = \pi(R^*)^2 L_d^* - \frac{2}{3}\pi(R^*)^3 = \pi[R(1+\rho) + \varepsilon_r]^2\left[\frac{A(1+\mu_d)}{2\pi R(1+\rho)} + \varepsilon_d\right] - \frac{2\pi[R(1+\rho)+\varepsilon_r]^3}{3}. \tag{S3}$$

The length at birth is derived from the length at division (defined by replacing $\mu_d$ with $\mu_b$) and by including the error $\alpha$ due to asymmetric mispositioning of the division septum. After division, contraction of the ring and turgor pressure deform the plane of division into a new hemispherical end while conserving the radius and cell volume [4], leading to an extra term $R(1+\rho)/3$:

$$L_b^* = \frac{A(1+\mu_b)(1+\alpha)}{4\pi R(1+\rho)} + \frac{R(1+\rho)}{3} + \varepsilon_b. \tag{S4}$$

We also verified that a use of a different fraction, e.g. $R(1+\rho)/2$, does not affect our results. Similar to above, we also derived expressions for the measured area and the measured volume at birth. These equations allow us to calculate all the quantities measured in the experiments, namely the 4 coefficients of variation (for radius, length, area and volume), and the slopes of the 3 division-birth plots (using length, area or volume as the geometrical feature used for size control).

**Coefficients of variation:** The coefficient of variation of X is defined as $\mathrm{CV}_X^2 = \sigma_X^2/\mathbb{E}[X]^2$, where $\mathbb{E}[X]$ denotes the expected value of the random variable *X*. As an example, we report here the calculation of the Coefficient of Variation (CV) of the measured length at division in the case of area size sensing, i.e. when:

$$L_d^* = \frac{A(1+\mu_d)}{2\pi R(1+\rho)} + \varepsilon_d.$$

Because of the small value of $\sigma_\rho$, the second-order approximation $(1+\rho)^{-m} \approx 1 - m\rho + \frac{m(m+1)}{2}\rho^2$ is used in all calculations. The definition of the variance is

$$\sigma_{L_d^*}^2 = \mathbb{E}[(L_d^*)^2] - \mathbb{E}[L_d^*]^2.$$

Each term is then calculated as follows:

$$\mathbb{E}[(L_d^*)^2] = \iiint_{-\infty}^{+\infty}\left[\frac{A(1+\mu_d)}{2\pi R(1+\rho)} + \varepsilon_d\right]^2 P(\mu_d)P(\rho)P(\varepsilon_d)d\mu_d d\rho d\varepsilon_d =$$

$$\approx \left(\frac{A}{2\pi R}\right)^2 \iint_{-\infty}^{+\infty}(1+2\mu_d+\mu_d^2)(1-2\rho+3\rho^2)P(\mu_d)P(\rho)d\mu_d d\rho + \int_{-\infty}^{+\infty}\varepsilon_d^2 P(\varepsilon_d)d\varepsilon_d$$

$$+ \iiint_{-\infty}^{+\infty}\frac{A(1+\mu_d)\varepsilon_d}{\pi R(1+\rho)}P(\mu_d)P(\rho)P(\varepsilon_d)d\mu_d d\rho d\varepsilon_d.$$

Since each Gaussian variable has zero mean value and retaining only lowest order terms, we have:

$$\mathbb{E}[(L_d^*)^2] = \left(\frac{A}{2\pi R}\right)^2 (1+\sigma_\mu^2+3\sigma_\rho^2)+\sigma_\varepsilon^2.$$

$$\mathbb{E}[L_d^*] \approx \iiint_{-\infty}^{+\infty} \left[\frac{A}{2\pi R}(1+\mu_d)(1-\rho+\rho^2)+\varepsilon_d\right] P(\mu_d)P(\rho)P(\varepsilon_d)d\mu_d d\rho d\varepsilon_d = \frac{A}{2\pi R}(1+\sigma_\rho^2).$$

$$\mathrm{CV}_{L_d^*}^2 = \frac{\sigma_{L_d^*}^2}{\mathbb{E}[L_d^*]^2} = \frac{\mathbb{E}[(L_d^*)^2]-(\mathbb{E}[L_d^*])^2}{(\mathbb{E}[L_d^*])^2} \approx \sigma_\mu^2+\sigma_\rho^2+\left(\frac{2\pi R}{A}\right)^2\sigma_\varepsilon^2.$$

The same procedure was applied to all the other geometrical quantities (area, volume, radius). Here, we report only the final result for the other quantities. These expressions were then used to fit the data of Fig. 2B and extract parameter values for the errors.

$$\mathrm{CV}_{R^*}^2 = \sigma_\rho^2+\frac{\sigma_\varepsilon^2}{R^2}.$$

$$\mathrm{CV}_{L_d^*}^2 \approx \sigma_\mu^2+\sigma_\rho^2+\left(\frac{2\pi R}{A}\right)^2\sigma_\varepsilon^2.$$

$$\mathrm{CV}_{A_d^*}^2 \approx \sigma_\mu^2+\left[\frac{1}{R^2}+\left(\frac{2\pi R}{A}\right)^2\right]\sigma_\varepsilon^2.$$

$$\mathrm{CV}_{V_d^*}^2 \approx \frac{A^2R^2\sigma_\mu^2+(A^2R^2+16\pi^2R^6-8\pi AR^4)\sigma_\rho^2+4(A^2+5\pi^2R^4-4\pi AR^2)\sigma_\varepsilon^2}{A^2R^2+\frac{16}{9}\pi^2R^6-\frac{8}{3}\pi AR^4}$$

We also investigated whether the square of the surface area CV is smaller than the square of the length CV:

$$\mathrm{CV}_{A_d^*}^2-\mathrm{CV}_{L_d^*}^2 \approx \frac{\sigma_\varepsilon^2}{R^2}-\sigma_\rho^2.$$

In particular, we studied the sign of their difference. The right hand-side of the equation describes the linear relationship of the difference between the two CVs as a function of $\sigma_\varepsilon^2$. The negative intercept $-\sigma_\rho^2$ indicates that this difference can be negative if the error is sufficiently small. In particular, in order to have $\mathrm{CV}_{A_d^*}^2 - \mathrm{CV}_{L_d^*}^2 < 0$ we must have $\sigma_\varepsilon < R\sigma_\rho$ ,i.e. the error must not be bigger than the natural absolute variability of the radius (see Fig. 2B in the main text).

**Slopes:** We calculated the slopes for the plot of length at division ($L_d^*$) *vs* length at birth ($L_b^*$). The slope of the linear regression of a set of pairs $(x_i, y_i)$, $i = 1, \ldots, N$ is

$$\mathrm{slope}(y,x) = \frac{N\sum x_i y_i - \sum x_i \sum y_i}{N\sum x_i^2 - (\sum x_i)^2},$$

which, for large *N* and for our quantities ($L_d^*$ and $L_b^*$), can be rewritten as follows:

$$\mathrm{slope}(L_d^*, L_b^*) = \frac{\mathbb{E}[L_b^* L_d^*]-\mathbb{E}[L_b^*]\mathbb{E}[L_d^*]}{\mathbb{E}[(L_b^*)^2]-\mathbb{E}[L_b^*]^2} = \frac{\mathrm{cov}(L_d^*, L_b^*)}{\mathrm{var}(L_b^*)}.$$

This expression explains why for the error in size sensing ($\sigma_\mu$), despite stretching along both the x- and y-axes, it is only the x-axis stretch that affects the division-birth slope. The above expectation values can be calculated using the same procedure adopted for the calculation of the CVs described above. With this analytical procedure, we derived Eq. 2 of the main text. As a verification of the approximations used, we also ran numerical simulations. In particular, we implemented a computational model which, by using Eqs. S1 and S4 and by simulating n=1000 cells, reproduced the same values of the analytical expressions (see Fig. S3A).

**Equations for models based on length sensing and on volume sensing for size control:** We report here the expressions of the initial quantities used to derive the CVs at division, and the division-birth slope, for the cases of length sensing and volume sensing, respectively (see above for the case of surface area sensing).

Length sensing:

$$L_d^* = L(1+\mu_d) + \varepsilon_d.$$

$$A_d^* = 2\pi[R(1+\rho) + \varepsilon_r][L(1+\mu_d) + \varepsilon_d].$$

$$V_d^* = \pi[R(1+\rho) + \varepsilon_r]^2[L(1+\mu_d) + \varepsilon_d] - \frac{2\pi[R(1+\rho) + \varepsilon_r]^3}{3}.$$

$$\mathrm{CV}^2_{L_d^*} \approx \sigma_\mu^2 + \frac{\sigma_\varepsilon^2}{L^2}.$$

$$\mathrm{CV}^2_{A_d^*} \approx \sigma_\mu^2 + \sigma_\rho^2 + \left(\frac{1}{R^2} + \frac{1}{L^2}\right)\sigma_\varepsilon^2.$$

$$\mathrm{CV}^2_{V_d^*} \approx \frac{9L^2\sigma_\mu^2 + 36(L-R)^2\sigma_\rho^2 + 9\left[1 + \frac{4}{R^2}(L-R)^2\right]\sigma_\varepsilon^2}{(3L-2R)^2}$$

$$\mathrm{slope}(L_d^*, L_b^*) = 0, \text{ always.}$$

Volume sensing:

$$L_d^* = \frac{V(1+\mu_d)}{\pi R^2(1+\rho)^2} + \frac{2R(1+\rho)}{3} + \varepsilon_d.$$

$$A_d^* = 2\pi[R(1+\rho) + \varepsilon_r]\left[\frac{V(1+\mu_d)}{\pi R^2(1+\rho)^2} + \frac{2R(1+\rho)}{3} + \varepsilon_d\right].$$

$$V_d^* = \pi[R(1+\rho) + \varepsilon_r]^2\left[\frac{V(1+\mu_d)}{\pi R^2(1+\rho)^2} + \frac{2R(1+\rho)}{3} + \varepsilon_d\right] - \frac{2\pi[R(1+\rho) + \varepsilon_r]^3}{3}.$$

$$\mathrm{CV}^2_{L_d^*} \approx \frac{9V^2\sigma_\mu^2 + 4(3V - \pi R^3)^2\sigma_\rho^2 + 9\pi^2R^4\sigma_\varepsilon^2}{(3V + 2\pi R^3)^2}.$$

$$\mathrm{CV}^2_{A_d^*} \approx \frac{9V^2\sigma_\mu^2 + (3V - 4\pi R^3)^2\sigma_\rho^2 + \frac{1}{R^2}[(3V + 2\pi R^3)^2 + 9\pi^2R^6]\sigma_\varepsilon^2}{(3V + 2\pi R^3)^2}.$$

$$\mathrm{CV}^2_{V_d^*} \approx \sigma_\mu^2 + \left[\frac{\pi^2R^4}{V^2} + \left(\frac{2}{R} - \frac{2\pi R^2}{3V}\right)^2\right]\sigma_\varepsilon^2$$

$$\mathrm{slope}(L_d^*, L_b^*) \approx \frac{2\left[\left(\frac{V}{\pi R^2}\right)^2 + \frac{2R^2}{9} - \frac{V}{\pi R}\right]\sigma_\rho^2}{\left(\frac{V}{\pi R^2} - \frac{2R}{3}\right)^2\sigma_\rho^2 + \left(\frac{V}{2\pi R^2}\right)^2\sigma_\mu^2 + \left(\frac{V}{2\pi R^2} + \frac{R}{3}\right)^2\sigma_\alpha^2 + \sigma_\varepsilon^2}.$$

This last expression was used to calculate the slope for the *cdr2*Δ mutant (Fig. 4D) and for *E. coli*. In the former case, we used the same parameters values as for the wild-type, with a mean division length of 17 µm, whereas in the latter case we used parameters according to the data in the available literature [5]. In particular, we used the geometrical features of this bacterium ($R = 0.55$ µm, $V = 3.77$ µm$^3$) and we estimated the natural variability $\sigma_\rho \approx 3.5\%$ from the value of the CV of the cell width in different growing media. To show the robustness of the result, we perturbed each noise up to ±2% and checked the distribution of the obtained slopes (Fig. S6B).

**Variability of the real radius in a subset of cells selected by the measured radius:** In Fig. 3B of the main text, we showed how the division-birth length slope reduces when the natural variability of the cell radius is reduced. To enact this strategy, we selected a subset of cells that have reduced variability. In particular we chose cells whose measured radius fell in the range $R \pm w$ (i.e. mean value ± $w$). In order to use Eq. 2 to calculate the predicted value of the division-birth slope for this subset of cells, we first needed to know the natural variability of this subset, which depends also on the experimental measurement error. We already know that

$$R^* = R(1+\rho) + \varepsilon,$$

where R represents the average cell radius over the entire population. Suppose we have a cell with a given real radius $R_{\text{real}}$. First, we want to know the probability that the measured radius of this cell (i.e. $R_{\text{real}} + \varepsilon$) falls in $I_w = (R - w, R + w)$. This question is the equivalent of asking the probability that $\varepsilon$ belongs to the interval $(R - R_{\text{real}} - w, R - R_{\text{real}} + w)$. Because $\rho$ and $\varepsilon$ are independent random variables, we can just multiply this probability by the probability for a cell to have that real radius $R_{\text{real}}$, i.e. $\text{Prob}[R(1+\rho) = R_{\text{real}}] = \text{Prob}[\rho R = R_{\text{real}} - R]$. Therefore, the probability that a cell of radius $R_{\text{real}}$ is selected:

$$p(R_{\text{real}}) = \frac{1}{N}\text{Prob}[\rho R = R_{\text{real}} - R]\text{Prob}[R - R_{\text{real}} - w < \varepsilon < R - R_{\text{real}} + w\ ],$$

where $N$ is the normalization factor. This expression gives the distribution of the radius of the cell we select in $I_w$. The CV of this distribution then gives us the required natural variability.

For simplicity, we introduced $Z = R_{\text{real}} - R$ and rewrote the probability of a cell with radius $R_{\text{real}}$ to fall in $I_w$ as follows:

$$p(Z = z) = \frac{1}{N}\text{Prob}[\rho R = z]\text{Prob}[-w < \varepsilon + z < +w\ ] = \frac{1}{N}\frac{1}{R\sigma_\rho\sqrt{2\pi}}e^{-\frac{z^2}{2R^2{\sigma_\rho}^2}}\frac{1}{\sigma_\varepsilon\sqrt{2\pi}}\int_{-w}^{w} e^{-\frac{(q+z)^2}{2{\sigma_\varepsilon}^2}}dq$$

where the normalization factor is $N = \text{erf}\left(\frac{w}{\sqrt{2(R^2\sigma_\rho^2+\sigma_\varepsilon^2)}}\right)$.

By construction $\text{Var}[R_{\text{real}}]$=$\text{Var}[Z]$, which is equal to $\mathbb{E}[Z^2]$ because Z has zero mean value. This led to

$$\text{Var}[R_{\text{real}}] = R^2\sigma_\rho^2 - \frac{2wR^4\sigma_\rho^4}{R^2\sigma_\rho^2+\sigma_\varepsilon^2}\frac{e^{-\frac{w^2}{2(R^2\sigma_\rho^2+\sigma_\varepsilon^2)}}}{\sqrt{2\pi(R^2\sigma_\rho^2+\sigma_\varepsilon^2)}}\text{erf}\left(\frac{w}{\sqrt{2(R^2\sigma_\rho^2+\sigma_\varepsilon^2)}}\right)^{-1}.$$

The real natural variability is then the CV of $R_{\mathrm{real}}$, i.e.

$$\text{Nat. var.}\,(w) = \sigma_\rho \sqrt{1 - \frac{2R^2\sigma_\rho^2}{R^2\sigma_\rho^2+\sigma_\varepsilon^2}\frac{w}{\sqrt{2\pi\left(R^2\sigma_\rho^2+\sigma_\varepsilon^2\right)}}\frac{e^{-\frac{w^2}{2\left(R^2\sigma_\rho^2+\sigma_\varepsilon^2\right)}}}{\mathrm{erf}\left(\frac{w}{\sqrt{2\left(R^2\sigma_\rho^2+\sigma_\varepsilon^2\right)}}\right)}}. \tag{S5}$$

It is worth noticing that, because of the error, the accessible lower bound (in the limit $w \to 0$) of the natural variability is not zero, but rather $\text{Nat. var.}\,(w = 0) = \sigma_\rho \sqrt{\frac{\sigma_\varepsilon^2}{R^2\sigma_\rho^2+\sigma_\varepsilon^2}}$. Equation S5 gives the curve of Fig. S3B and was used to calculate the natural variability of the experimental data in Fig. 3B of the main text.

**Equations for the pure adder model**

A similar approach was used also for adder control with incremental size ΔV defined by the volume. We report here only the equations for length at birth and length at division. Because several generations are needed to converge to the theoretical size, we implemented only the numerical version of the model. The simulation was run by starting with a length at birth equal to $\Delta\mathrm{V}/\,\pi R^2$ and iterating 50 cell cycles. Analyses were performed on the cell size at the last cell cycle. Superscript [*n*] denotes a quantity at the *n*-th cell cycle.

$$L_b^{*[n]} = \frac{L_d^{[n-1]}(1+\alpha)}{2} + \frac{R(1+\rho)}{3} + \varepsilon_b.$$

$$L_d^{*[n]} = L_b^{[n]} + \frac{\Delta\mathrm{V}(1+\mu_d)}{\pi R^2(1+\rho)^2} + \varepsilon_d.$$

From the numerical simulation of 3000 independent cells, we derived the value of the slope $L_d^*$ *vs* $L_b^*$.

### SUPPLEMENTARY FIGURES

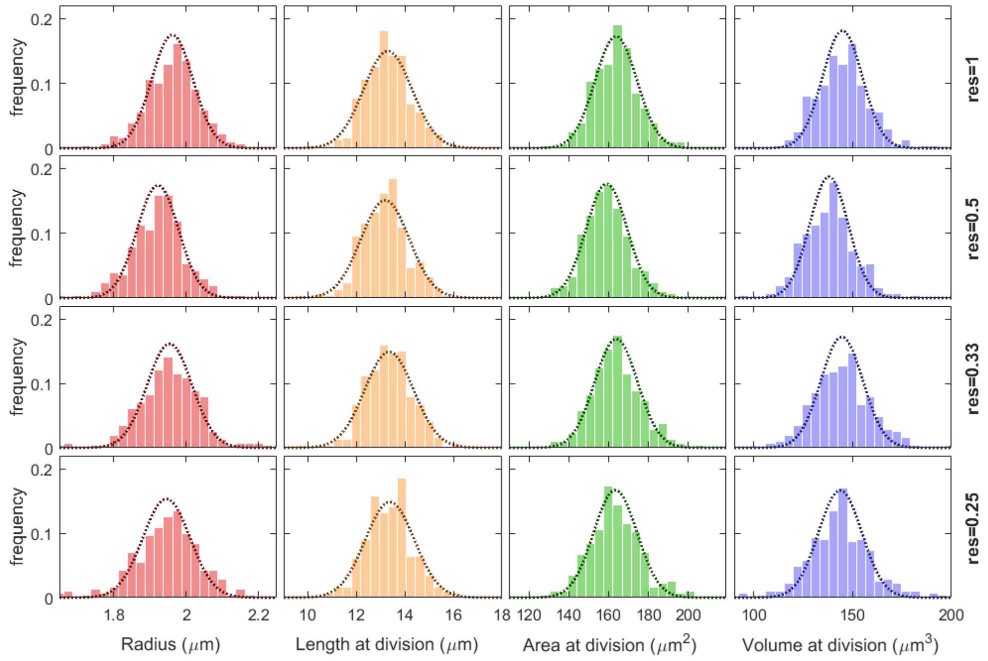


**Figure S1: Estimation of parameters.** Fitted distributions of the different geometrical quantities of cells at division for each resolution (Fig. 2A-B). Histogram bars: experimental data; dotted line: Gaussian distribution with variance obtained using the model equations.

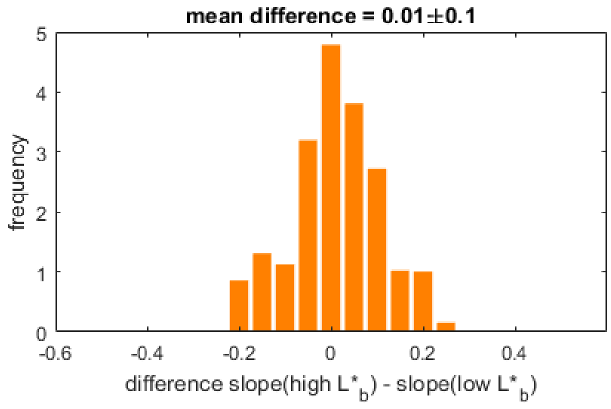


**Figure S2: Apparently imperfect sizer behaviour.** Slope of plot in Fig. 1A might be due to the presence of two regimes, i.e. a sizer regime (slope close to 0) at low birth lengths and an adder regime (slope close to 1) at high birth lengths (as in *pom1*Δ, see Fig. 4B in main text and [6]). To exclude this possibility, we split the cells into low and high birth lengths according to a threshold and calculated the slopes in the two regimes. We varied the value of the threshold and analysed the distribution of the difference between "slope at high $L_b$" and "slope at low $L_b$" (bar plot). Since the mean value of this distribution, as indicated in the figure, is very close to zero, we conclude that the cells do not show two-regime behaviour.

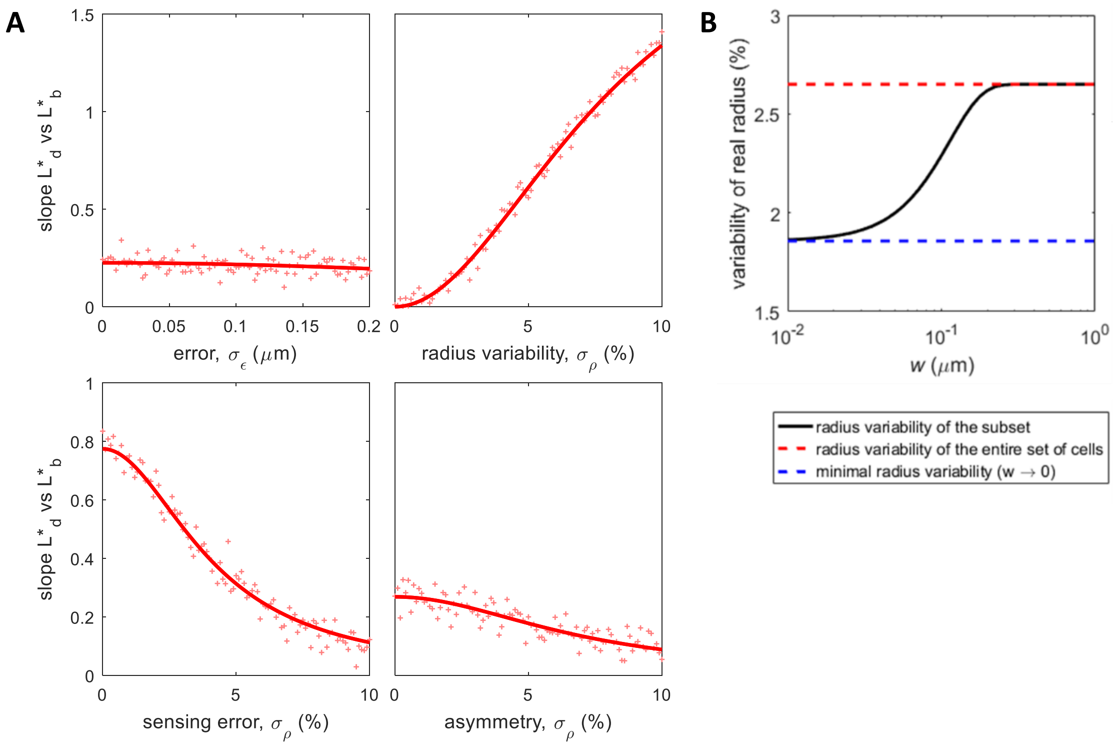


**Figure S3: Theoretical division-birth slopes for a pure area-based sizer control and effect of the noise sources.** (**A**) Effect of experimental error ($\sigma_\varepsilon$), radius variability ($\sigma_\rho$), sensing error ($\sigma_\mu$), asymmetry ($\sigma_\alpha$) on the division-birth slopes. Lines: analytical equations for division-birth slope ($L^*_d$ vs $L^*_b$), where subscript *d* refers to division and subscript *b* to birth, assuming underlying area-based size sensing. "+": numerical simulation (n=1000 cells). Except for the x-axis values, all other parameters are as in Table S1. (**B**) Variability of the real radius for the subset of cells with measured radius $R^*$ in the range $R \pm w$ (see Eq. S5). This analysis is used to derive the *x*-values in Fig. 3B.

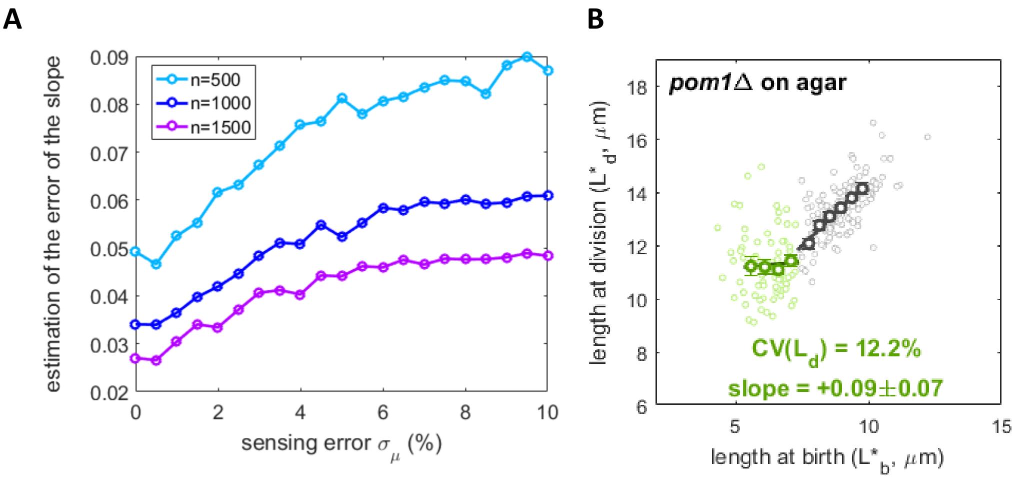


**Figure S4: Effect of sensing error $\sigma_\mu$ on division-birth slope uncertainty and repeated *pom1*Δ experiment.** (**A**) Estimation of the uncertainty of the division-birth slope as standard deviation over 1000 independent numerical simulations of n cells. To maintain the same error, a larger number of cells (n) must be imaged. (**B**) Size homeostasis plot for *pom1*Δ. Binned data, with mean value ± standard error, also shown (dark circles), together with best fit line. Slope of the second regime: +0.97±0.07. (FC2063, n=402).

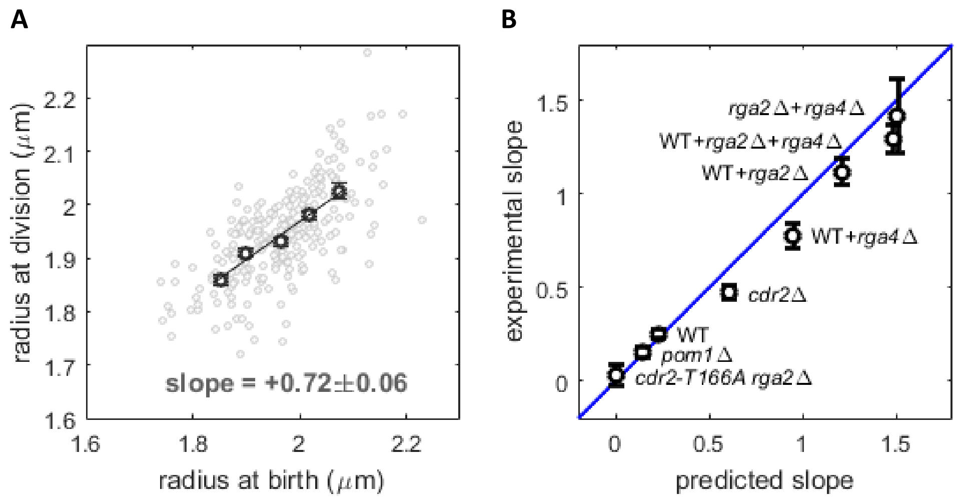


**Figure S5: Model with radius change during the cell cycle.** (**A**) Experimental data from wild-type cells (FC15, n=256). Binned data, with mean value ± standard error, also shown (dark circles), together with best fit line with slope $q$=0.72 and intercept 0.54, which is in good agreement with our theoretical value of $R(1-q) = 0.55$. (**B**) Comparison between experimental and predicted slope by using the model with radius changes. Data points show the mean value ± standard error. See also Fig. 4E.

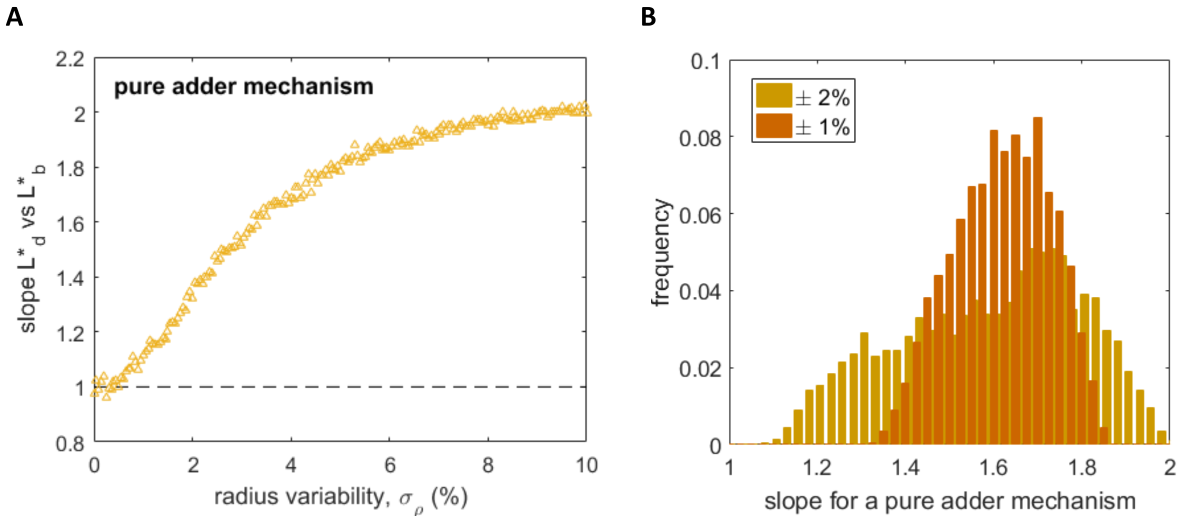


**Figure S6: Paradoxes in size behaviour.** (**A**) Effect of radius variability on the division-birth length slope in case of a pure adder mechanism based on volume control. Other parameters values: noise as for the other model predictions ($\sigma_\varepsilon = 0.04$ µm, $\sigma_\mu = 6.5\%$, $\sigma_\alpha = 3.2\%$), cell size $R = 0.55$ µm, $V = 3.77$ µm$^3$ [5]. (**B**) Distribution of slopes when additively increasing/decreasing each standard deviation by up to 1% (or 2%) for all parameters of panel A, with fixed $\sigma_\rho = 3.5\%$.

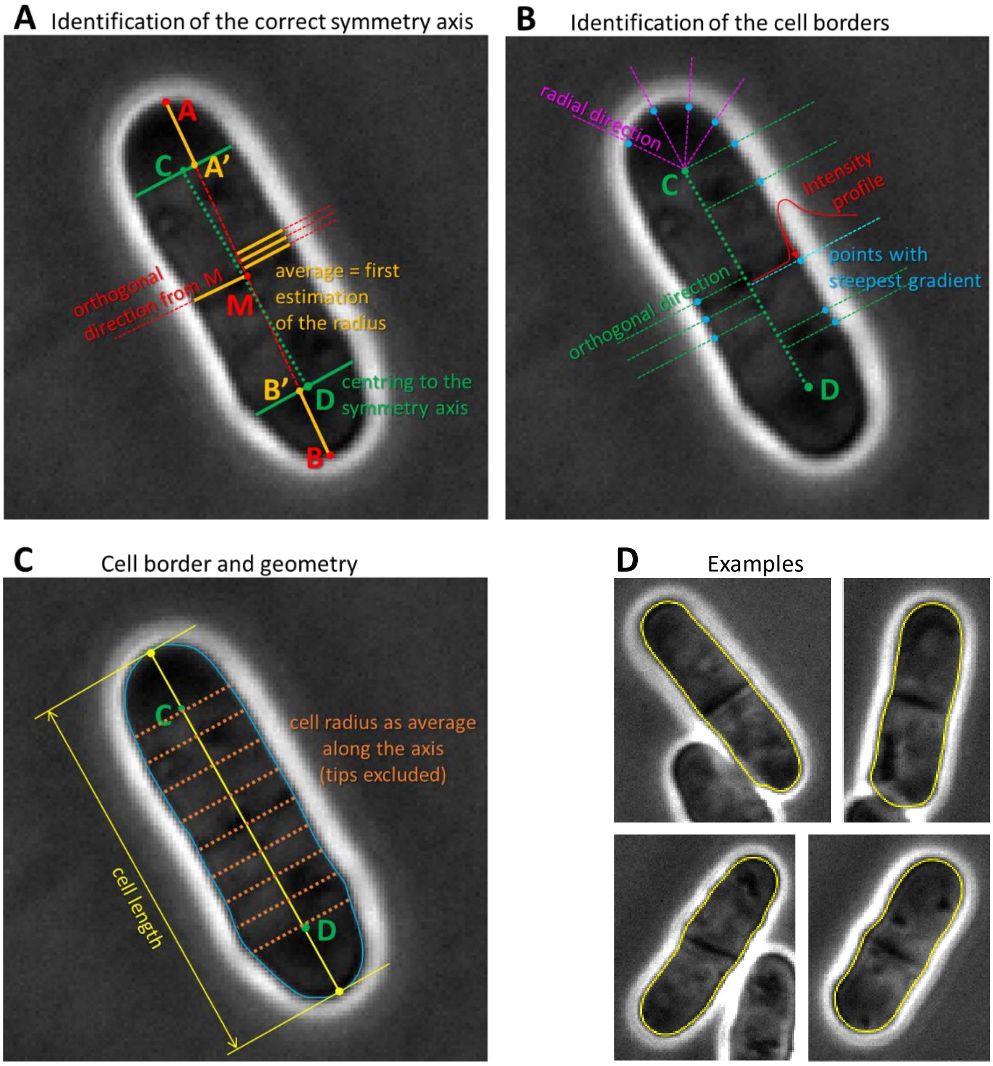


**Figure S7: Semi-automated segmentation algorithm**. (**A-C**) Sequence of operations performed to segment the cell border (see SI text). (**D**) Examples of segmented cells.

### SUPPLEMENTARY TABLES

**TABLE S1: Parameter values from fitting the CVs for each type of geometric size sensing** (Fig. 2B and Fig. 3D-E).

| *parameter* | $\sigma_\rho$ | $\sigma_\mu$ | $\sigma_\varepsilon$ (µm) at different resolutions | | |
|---|---|---|---|---|---|
| Length sensing | 1.7 % | 6.8 % | 0.036 | res.=1 | 0.0635 µm/pixel |
| | | | 0.043 | res.=0.5 | 0.127 µm/pixel |
| | | | 0.059 | res.=0.33 | 0.190 µm/pixel |
| | | | 0.072 | res.=0.25 | 0.254 µm/pixel |
| Area sensing | 2.7 % | 6.5 % | 0.045 | res.=1 | 0.0635 µm/pixel |
| | | | 0.052 | res.=0.5 | 0.127 µm/pixel |
| | | | 0.067 | res.=0.33 | 0.190 µm/pixel |
| | | | 0.078 | res.=0.25 | 0.254 µm/pixel |
| Volume sensing | 2.0 % | 6.8% | 0.058 | res.=1 | 0.0635 µm/pixel |
| | | | 0.062 | res.=0.5 | 0.127 µm/pixel |
| | | | 0.075 | res.=0.33 | 0.190 µm/pixel |
| | | | 0.085 | res.=0.25 | 0.254 µm/pixel |

**TABLE S2: Strains used in this study.**

| *S. pombe* strain | SOURCE |
|---|---|
| FC15: $h^-$ *WT (972)* | Lab collection |
| FC2947: $h^-$ *rga2::ura4$^+$ ade6- leu1-32 ura4-D18* | Lab collection |
| FC1901: $h^-$ *rga4::ura4$^+$ leu1-32 ura4-D18* | Lab collection |
| FC2063: $h^-$ *pom1::natMX4 ade6- leu1-32 ura4-D18* | Lab collection |
| FC3161: $h^+$ *cdr2::kanMX leu1-32* | Lab collection |
| FC3218: $h^-$ *cdr2-T166A rga2::ura4$^+$* | Lab collection |

### SUPPLEMENTARY REFERENCES


1. Facchetti, G., Knapp, B., Flor-Parra, I., Chang, F., and Howard, M. (2019). Reprogramming Cdr2-Dependent Geometry-Based Cell Size Control in Fission Yeast. Current Biology.
2. Van Valen, D.A., Kudo, T., Lane, K.M., Macklin, D.N., Quach, N.T., DeFelice, M.M., Maayan, I., Tanouchi, Y., Ashley, E.A., and Covert, M.W. (2016). Deep learning automates the quantitative analysis of individual cells in live-cell imaging experiments. PLoS computational biology *12*, e1005177.
3. Ursell, T., Lee, T.K., Shiomi, D., Shi, H., Tropini, C., Monds, R.D., Colavin, A., Billings, G., Bhaya-Grossman, I., and Broxton, M. (2017). Rapid, precise quantification of bacterial cellular dimensions across a genomic-scale knockout library. BMC biology *15*, 17.
4. Baumgärtner, S., and Tolić-Nørrelykke, I.M. (2009). Growth pattern of single fission yeast cells is bilinear and depends on temperature and DNA synthesis. Biophysical journal *96*, 4336-4347.
5. Taheri-Araghi, S., Bradde, S., Sauls, J.T., Hill, N.S., Levin, P.A., Paulsson, J., Vergassola, M., and Jun, S. (2015). Cell-size control and homeostasis in bacteria. Current Biology *25*, 385-391.
6. Wood, E., and Nurse, P. (2013). Pom1 and cell size homeostasis in fission yeast. Cell cycle *12*, 3417-3425.
